# Non-covalently-associated peptides are observed during liquid chromatography-mass spectrometry and affect crosslink analyses

**DOI:** 10.1101/502351

**Authors:** Sven H. Giese, Adam Belsom, Ludwig Sinn, Lutz Fischer, Juri Rappsilber

**Affiliations:** Bioanalytics, Institute of Biotechnology, Technische Universität Berlin, 13355 Berlin, Germany; Wellcome Centre for Cell Biology, School of Biological Sciences, University of Edinburgh, Edinburgh EH93BF, United Kingdom

## Abstract

Crosslinking mass spectrometry draws structural information from covalently-linked peptide pairs. When these links do not match to previous structural models, they may indicate changes in protein conformation. Unfortunately, such links can also be the result of experimental error or artefacts. Here, we describe the observation of non-covalently-associated peptides during liquid chromatography-mass spectrometry analysis, which can easily be misidentified as crosslinked. Strikingly, they often mismatch to the protein structure. Non-covalently-associated peptides presumably form during ionization and can be distinguished from crosslinked peptides by observing co-elution of the corresponding linear peptides in MS1, as well as the presence of the individual (intact) peptide fragments in MS2 spectra. To suppress non-covalent peptide formations increasingly disruptive ionization settings can be used, such as in-source fragmentation.

---

The preservation of non-covalent associations in electrospray ionization (ESI) has been widely used in the field of native mass spectrometry to study protein interactions. Major achievements of native mass spectrometry include analyzing the topology and stoichiometry of multi-protein complexes and the binding of small molecules to proteins.^1–3^ The key premise of the field is that the observed non-covalent interactions in the gas phase are based on biologically relevant interactions in the aqueous phase.^4^

Another mass spectrometric field that investigates (non-)covalent interactions of proteins is crosslinking mass spectrometry (CLMS).^5–7^ Here, spatially close amino acid residues in native proteins are covalently linked. This preserves spatial information throughout the subsequent non-native analytical process, comprising trypsin digestion of the proteins into peptides and their chromatographic separation for mass spectrometric detection. A key premise of this field is that the observed peptide interactions in the gas phase are exclusively based on covalent links. Note that for synthetic peptides, gas-phase peptide-peptide complexes have been observed recently^8^, suggesting that not only proteins but also peptides can remain associated during mass spectrometric analysis.

In theory, one can construct peptide pairs where mass information alone cannot differentiate between covalent linkage and non-covalent association. A peptide pair can reach the same mass either by crosslinking or by non-covalent association if one of the two peptides carries a loop-link, i.e. the frequent case of a crosslinker reacting with two amino acid residues so near in sequence that they fall into a tryptic peptide (Fig. S1). The concept of mass-equivalence between crosslinked and not crosslinked peptides has been exploited during data analysis, when using standard proteomics software for the analysis of crosslinked peptides, including Mascot^9^ to identify crosslinked peptides^10^ and quantitation software^11,12^. If such non-covalent associations physically arise, current crosslink analysis could be fooled into misidentifying analytical artifacts as spatial information.

We observed surprising differences when comparing the identified crosslinks using data acquired on two different mass spectrometers: a hybrid linear ion trap-Orbitrap mass spectrometer (LTQ Orbitrap Velos, Thermo Fisher Scientific) and a hybrid quadrupole-Orbitrap mass spectrometer (Q Exactive, Thermo Fisher Scientific). This led us to investigate the formation of non-covalent peptide associations with and without crosslinking. We analyzed crosslinked human serum albumin (HSA). Using only the monomeric protein band obtained from an SDS-PAGE allowed identified links to be validated against an available three-dimensional structural model as “ground truth” to reveal suspicious peptide pairs for detailed interrogation. We then extended this data analysis to a four-protein mix without employing crosslinking to test if the non-covalent association is crosslinker specific.

## Materials & Methods

### Data Acquisition

#### HSA acquisition and sample preparation

Human blood serum (20 μg aliquots, 1 μg/μL) was crosslinked using crosslinker-to-protein, weight-to-weight (w/w) ratios of 1:1 and 2:1. Aliquots of human serum diluted with crosslinking buffer (20 mM HEPES-OH, 20 mM NaCl, 5 mM MgCl2, pH 7.8) were incubated with sulfosuccinimidyl 4,4’-azipentanoate (sulfo-SDA) (Thermo Scientific Pierce, Rockford, IL), in a reaction volume of 30 μL for 1 h at room temperature. The diazirine group was then photo-activated by UV irradiation, for either 10, 20, 40 or 60 minutes using a UVP CL-1000 UV Cross-linker (UVP Inc.). Crosslinked samples were separated using gel electrophoresis, with bands corresponding to monomeric HSA excised then reduced with dithiothreitol, alkylated with iodoacetamide, and digested using trypsin following standard protocols.^10^ Peptides were then desalted using C18 StageTips^13^ and eluted with 80% acetonitrile, 20% water and 0.1% TFA.

Peptides were analyzed on either a hybrid linear ion trap/Orbitrap mass spectrometer (LTQ Orbitrap Velos, Thermo Fisher Scientific) or a hybrid quadrupole/Orbitrap mass spectrometer (Q Exactive, Thermo Fisher Scientific). In both cases, peptides were loaded directly onto a spray analytical column (75 μm inner diameter, 8 μm opening, 250 mm length; New Objectives, Woburn, MA) packed with C18 material (ReproSil-Pur C18-AQ 3 μm; Dr. Maisch GmbH, Ammerbuch-Entringen, Germany) using an air pressure pump (Proxeon Biosystems)^14^.

#### Orbitrap Velos analysis

Mobile phase A consisted of water and 0.1% formic acid. Peptides were loaded using a flow rate of 0.7 μl/min and eluted at 0.3 μl/min, using a gradient with a 1-minute linear increase of mobile phase B (acetonitrile and 0.1% v/v formic acid) from 1% to 9%, increasing linearly to 35% B in 169 minutes, with a subsequent linear increase to 85% B over 5 minutes. Eluted peptides were sprayed directly into the hybrid linear ion trap-Orbitrap mass spectrometer. MS data were acquired in the data-dependent mode, detecting in the Orbitrap at 100,000 resolution. The eight most intense ions in the MS spectrum for each acquisition cycle, with a precursor charge state of 3+ or greater, were isolated with a m/z window of 2 Th and fragmented in the linear ion trap with collision-induced dissociation (CID) at a normalized collision energy of 35. Subsequent (MS2) fragmentation spectra were then recorded in the Orbitrap at a resolution of 7500. Dynamic exclusion was enabled with single repeat count for 90 seconds.

#### Q Exactive analysis

Mobile phase A consisted of water and 0.1% formic acid. Mobile phase B consisted of 80% v/v acetonitrile and 0.1% formic acid. Peptides were loaded at a flow rate of 0.5 μl/min and eluted at 0.2 μl/min, using a gradient increasing linearly from 2% B to 40% B in 169 minutes, with a subsequent linear increase to 95% B over 11 minutes. Eluted peptides were sprayed directly into the hybrid quadrupole-Orbitrap mass spectrometer. MS data (400–1600 m/z) were acquired in the data-dependent mode, detecting in the Orbitrap at 60,000 resolution. The ten most intense ions in the MS spectrum, with a precursor charge state of 3+ or greater, were isolated with a m/z window of 2 Th and fragmented by Higher Energy Collision-Induced Dissociation (HCD) at a normalized collision energy of 28. Subsequent (MS2) fragmentation spectra were recorded in the Orbitrap at a resolution of 30,000. Dynamic exclusion was enabled with single repeat count for 60 seconds.

#### In-source collision induced dissociation acquisitions

HSA, equine myoglobin, ovotransferrin from chicken (all from Sigma-Aldrich, St. Louis, MO) and creatine kinase from rabbit (Roche, Basel, Switzerland) were dissolved in 8 M urea with 50 mM ammonium bicarbonate to a concentration of 2 mg/mL each. The proteins were reduced by adding dithiothreitol at 2.5 mM followed by an incubation for 30 minutes at 20 °C. Subsequently, the samples were derivatized using iodoacetamide at 5 mM concentration for 20 minutes in the dark at 20 °C. The samples were diluted 1:5 with 50 mM ammonium bicarbonate and digested with trypsin (Pierce Biotechnology, Waltham, MA) at a protease-to-protein ratio of 1:100 (w/w) during a 16-hour incubation period at 37 °C. Then, the digestion was stopped by adding 10% TFA at a concentration of 0.5%. The digests were cleaned up using the StageTip protocol.^13^ The samples were eluted from C18-phase, partially evaporated using a vacuum concentrator and resuspended in mobile phase A (0.1% formic acid). Two micrograms of tryptic digests were loaded directly onto a 50 cm EASY-Spray column (Thermo Fisher) packed with C18-stationary phase and equilibrated to 2% of mobile phase B (80% acetonitrile, 0.1 % formic acid) running at a flow of 0.3 μl/min. Peptides were eluted by increasing mobile phase B content from 2 to 37.5% over 120 minutes, followed by ramping to 45% and to 95% within 5 minutes each. After a washing period of 5 minutes, the column was reequilibrated to 2% B. The eluting peptides were sprayed into a Q Exactive High-field (HF) Hybrid Quadrupole-Orbitrap Mass Spectrometer (Thermo Fisher Scientific, Bremen, Germany). The mass spectrometric measurements in data-dependent mode were acquired as follows: a full scan from 400–1600 m/z with a resolution of 120,000 was recorded to find suitable peptide candidates which were subsequently quadrupole-isolated within a m/z window of 2 Th and fragmented by HCD at a normalized collision energy of 28, with fragmentation spectra recorded in the Orbitrap at a resolution of 30,000. Precursors with charge-states from 3 to 6 were selected for isolation. Dynamic exclusion was set to 15 seconds. Each cycle allowed up to ten peptides to be fragmented before a new full scan was triggered. The effect of in-source collisional activation (ISCID) on the formation of non-covalently bound peptides was investigated by setting voltages from 0 to 20 eV in 5 eV increments for each individual run. Each value tested was probed in shuffled triplicates.

### Data Processing

Raw files for crosslinking searches were processed using MaxQuant^15^ (v. 1.6.1.0) to benefit from the implemented precursor m/z and charge correction. Resulting peak files in APL format were used to identify peptides in Xi^16^ (v. 1.6.739). The database search with Xi used the following parameters: MS tolerance, 6 ppm; MS2 tolerance, 15 ppm; missed cleavages, 3; enzyme, trypsin; fixed modifications, carbamidomethylation (cm, +57.02 Da); variable modifications, oxidation methionine (ox, +15.99 Da). For sulfo-SDA, the crosslinker mass 82.04 Da and the modifications SDA-loop (+82.04 Da) and SDA-hyd (+100.05 Da) were used.^17^ False discovery rate (FDR) estimation was done using xiFDR^18^ (v. 1.1.26.58), using either 5% link FDR (without boosting) or a 5% peptide spectrum match (PSM) FDR. The Euclidean crosslink distances within HSA were estimated from mapping the peptide sequences to the three-dimensional structure when possible (PDB: 1AO6^19^).

Searches for non-covalently associated peptides (NAP) in the absence of crosslinkers were also conducted using Xi with a feature to search for non-covalently associated peptides. FDR analysis was done at a 5% PSM level using the formula: FDR=(TD-DD)/TT,^18^ after removing all PSMs with a score less than 1. FDRs were then transformed to q-values, defined as the minimal FDR at which a PSM would pass the threshold.^20^

Linear peptide identifications from crosslinked acquisitions were done using MaxQuant. We added the above defined SDA-loop and SDA-hyd modifications to the configuration file and allowed up to 5 modifications on a peptide together with a maximum of 5 missed cleavages. Resulting peptide identifications were filtered at the default FDR of 1%. Non-crosslinked acquisitions were searched with default settings treating each replicate as different experiment in the experimental design.

RT profiles for a given m/z were extracted using the MS1 (peak picked) raw data after conversion to mzML using msconvert^21^. The post-processing was done in Python using pyOpenMS^22^. RT profiles were defined as intensity values for a given m/z for the monoisotopic peak and two isotope peaks. During the developed look-up strategy, the precursor m/z of the identified crosslinked peptide, the m/z of the alpha peptide and the m/z of the beta peptide were searched in the MS1 data. The precursor mass only matches the sum of the individual peptides in a non-covalently associated peptide if one of the two peptides is SDA-loop-modified. Therefore, the MS1 data was screened for m/z traces of the individual peptides with and without an added SDA-loop modification. Similarly, all charge states up to the precursor charge were used. The m/z trace with the largest number of peaks was eventually selected for each individual peptide. The m/z seeds were all treated similarly, in a RT window of 180 seconds the given m/z was searched with a 20-ppm tolerance. If the m/z was found, the intensity was extracted. Resulting RT profiles where smoothened by a moving average with 15 points. For further data processing and visualization, the RT profile with the most peaks (either monoisotopic, first or second isotope peak) was selected.

Statistical analysis and data processing was performed using Python and the scientific package SciPy^23^. Unless otherwise noted, we performed significant tests using one-sided Mann-Whitney-U-Tests with α=0.05 and continuity correction. We used the following encoding for p-values: ns – not significant, * ≤ 0.05, ** ≤ 0.01, *** ≤ 0.001. Along with the significance tests, we provided effect size estimates based on Cohen’s d^24^ with pooled standard deviations, which uses the following classification: small – |d|≥0.2, medium – |d|≥0.5 and large – |d|≥0.8.

The mass spectrometry raw files, peak lists, search engine results, MaxQuant parameter files and FASTA files have been deposited to the ProteomeXchange Consortium (http://proteomecentral.proteomexchange.org) via the PRIDE partner repository^25^ with the dataset identifier PXD010895.

### Results And Discussion

The results are divided into 4 parts: (1) describes the results from HSA crosslinked using sulfosuccinimidyl 4,4’-azipentanoate (sulfo-SDA) and then analyzed with a Q Exactive (QE) and LTQ Orbitrap Velos (Velos) mass spectrometer, (2) describes the MS2 properties of the detected long-distance links (LDL) with the QE and introduces the hypothesis of non-covalently associated peptides (NAP) enduring ESI, (3) summarizes intensity and retention time (RT) properties of the identified PSMs and (4) shows that non-covalently associated peptides also occur in the absence of crosslinking.

#### Instrument comparison reveals a high number of suspicious crosslinks in Q Exactive data

We started by comparing the results from crosslinking HSA with sulfo-SDA using two different mass spectrometers – a Velos and a QE. Crosslinked peptides were identified using Xi with subsequent FDR filtering using xiFDR at a 5% link-level FDR. To independently assess the quality of the results, we evaluated how the identified crosslinks matched to the available crystal structure of HSA. At 5% link FDR, we identified 449 (QE) and 240 (Velos) links, of which 430 and 231 could be mapped to the available sequence in the structural model, respectively. The distance distributions of the mapped crosslinks looked similar for links below 22 Å (Fig. 1a). However, for long-distances, the link distributions looked different. The QE data shows a much higher percentage of links exceeding the 25 Å cutoff, which is the empirically defined distance limit of SDA crosslinking^26^. This leads to 18% long-distance links (LDL) for the QE data compared to 2% for the Velos data (Fig. 1a inlet). Since the protein monomer band was analyzed, the possibility that the LDL were derived from crosslinked homooligomers can be largely neglected. One possibility is that the deeper analysis on the QE, which is faster and more sensitive than the Velos, detected a rare protein conformational state. However, a previous analysis of SDA-crosslinked HSA on the Velos yielded 500 identified links (5% link FDR), with comparatively few LDL (6%)^27^. Also, data on the much faster and more sensitive Fusion Lumos did not return in our hands such proportion of LDLs (data not shown). This suggests that the QE data does not cover conformational flexibility of the protein. Instead, the QE data appears to suffer from a systematic error that leads to many false identifications. Importantly, this bias only affects target sequences as it is not controlled by the FDR estimation. If these LDL were indeed based on false identifications, one could suspect that they were identified based on weak data and thus derived from low scoring PSMs. We therefore compared the highest scoring PSM for each link above and below 25 Å (Fig. 1b). Remarkably, the LDL showed an even higher average score than the within-distance links. This difference was small and not significant, but it was still surprising that the two classes had a similar score distribution. Next, we manually investigated LDL PSMs to identify characteristics that might lead to a mechanistic explanation of these links.

**Figure 1:**
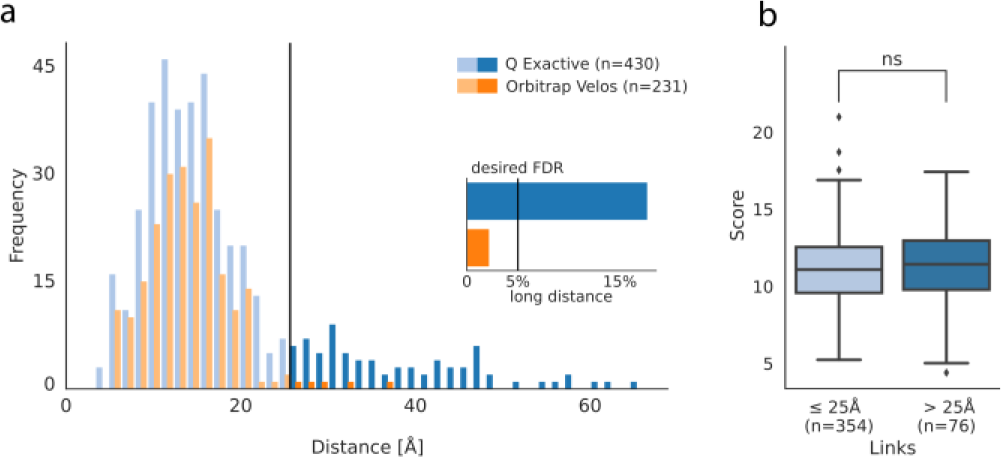
Quality control after crosslink identification at a 5% link FDR. (a) Results from crosslinking HSA with sulfo-SDA acquired on an Q Exactive and an LTQ Orbitrap Velos mass spectrometer. The line at 25 Å indicates the distance cutoff for links classified as long-distance. The inlet shows the fraction of long-distance links (LDL) in each dataset. (b) Score comparison between within-distance and LDL. LDL showed no significant (ns) deviation from the within-distance links (two-sided Mann-Whitney-U-test at α = 0.05).

#### Long-distance links lack support for being crosslinked

After suspecting a systematic identification error in QE data, we manually inspected annotated LDL spectra. We noticed that many spectra frequently contained unexplained fragment peaks of high intensity. For example, in the displayed spectrum (Fig. 2a, upper panel), most of the high intensity peaks are explained but not the base peak. This PSM was matched with a very low precursor error of 0.44 ppm and had a very good sequence coverage in general. However, while many of the linear fragments were identified, no crosslinked fragments were matched. While there is convincing evidence that the two identified peptides are correct, there is a lack of fragment evidence that these peptides were indeed crosslinked.

**Figure 2:**
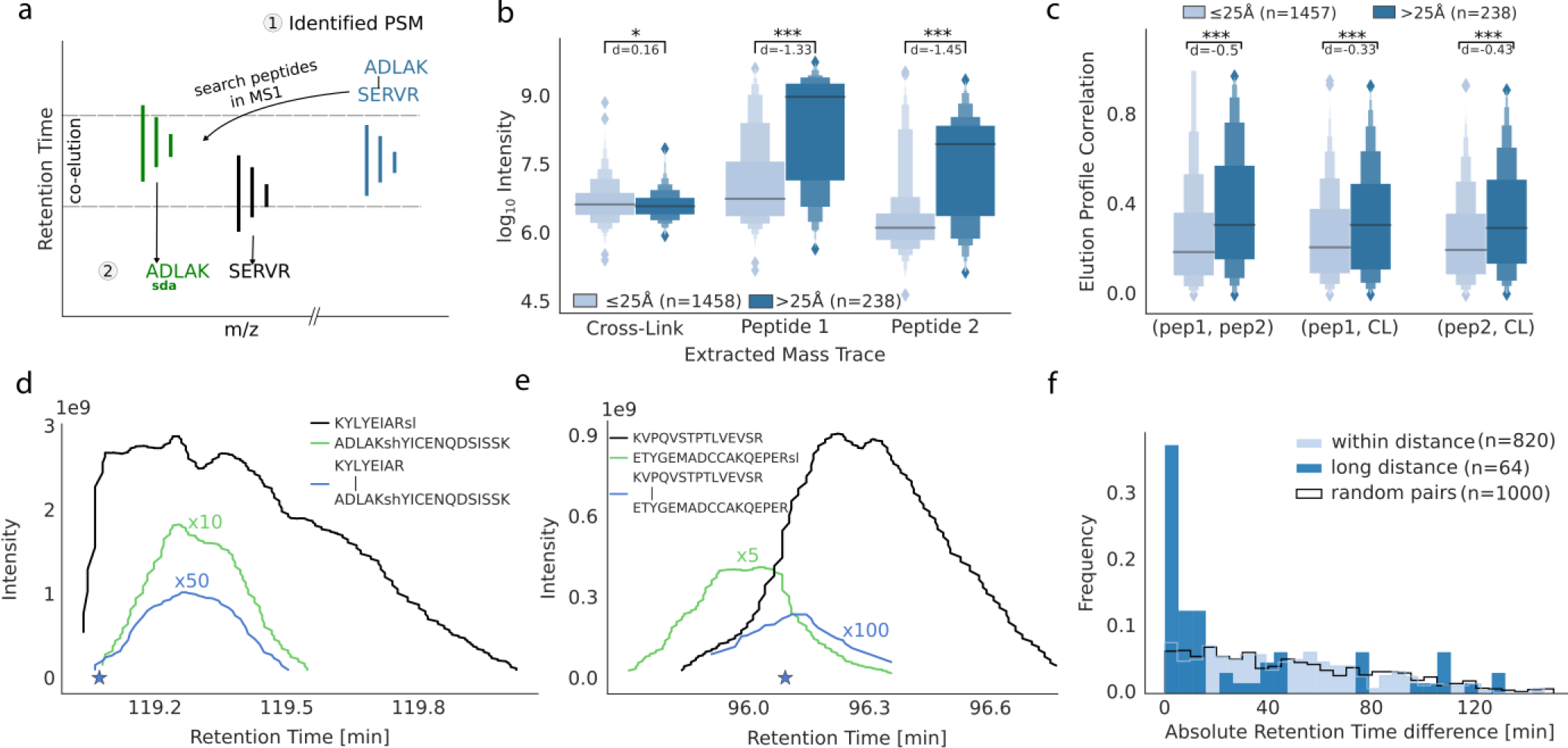
Spectral characteristic of non-covalently associated peptides. (a) Comparison of the same scan (scan 34887, raw file *V127_F*) searched with crosslink settings (upper panel) and searched with a non-covalent association setting (lower panel). (b) Comparison of the explained intensity in the MS2 spectrum from all PSMs that passed the 5% link-level FDR. (c) Comparison of crosslinker-containing fragments and linear fragments in the same set of PSMs as in (b). Number of observations for (b) and (c): ≤ 25 Å 2599 PSMs and > 25 Å 326 PSMs.

We tested our manual observations more systematically by comparing the explained intensity in the MS2 spectrum across all PSMs that passed the 5% link FDR (Fig. 2b). There is already a two-fold increase in the median explained intensity (EI) of the within-distance links (20% EI) and the LDL (10% EI). This trend is also supported by a significant MWU-test (one-sided, α=0.05) and a large Cohen’s d effect size (d=0.95). One possible explanation is that the spectra that yield LDL are simply of poor quality. This can happen when, for example, peptides of similar m/z were co-isolated, the precursor was of low intensity or when the peptide simply did not fragment or ionize very well. But as shown in Fig. 1b, the search engine scores of LDL were slightly higher than the scores from within-distance links. Therefore, poor spectral quality is not a likely reason for the large proportion of LDL. However, the number of matched crosslinked and linear fragments was significantly lower for the long-distance matches compared to the within-distance matches (Fig. 2c).

Recently, it has been proposed that SDA-formed bonds are very susceptible to MS cleavage when involving a carboxylic acid functional group^28^. In these cases, the annotated spectra would also show a low EI and a low number of crosslinked fragments with our search settings. However, it is unclear why such a reaction should preferentially lead to LDLs. Therefore, we hypothesized that the respective peptide pairs were not actually crosslinked but were non-covalently associated. Nevertheless, we investigated this in larger detail by following the approach of Iacobucci *et al*.^28^ and performed a cleavable crosslinker search on the Velos and the QE acquisitions (Fig. S2). A large portion of the identifications from the cleavable crosslinker search on the QE (38%) were long-distance links (presumably non-covalent peptide associations). However, the distribution of links that match the crystal structure revealed a preference for short distances thereby indicating that MS cleavage of the crosslinker can indeed be observed. So, our data support both as parallel processes MS-cleavable SDA links and non-covalent peptide complexes.

It would be interesting to investigate sequence determinants of non-covalent association. Unfortunately, the lack of ground truth and the low number of observations makes it difficult to investigate sequence specific features that lead to non-covalent peptide complexes. While crosslinks should preferentially fall below the distance cut-off, non-covalent peptide associations should distribute randomly across the distance histogram. Therefore, some links that match the crystal structure will also arise from non-covalent associations. Those links falling above the distance cut-off were too low in numbers for a statistical enrichment analysis.

#### Non-covalently associated peptides are of low intensity and arise when two individual peptides co-elute

As shown above, LDL frequently achieved high scores and there was good evidence based on the MS2 fragmentation that the peptides were correctly identified. Had the peptides paired non-covalently, this could either happen in solution or during the ESI process. In the latter case, one would expect the individual peptides to overlap in their chromatographic elution forming a non-covalent pair during their co-elution. In contrast, for crosslinked peptides one would not expect any systematic co-elution. Therefore, we investigated the elution of the individual peptides for all identifications (5% PSM FDR) following a look-up strategy that started from the MS2 trigger time of the crosslinked PSM (Fig. 3a, for details see material & methods).

**Figure 3:**
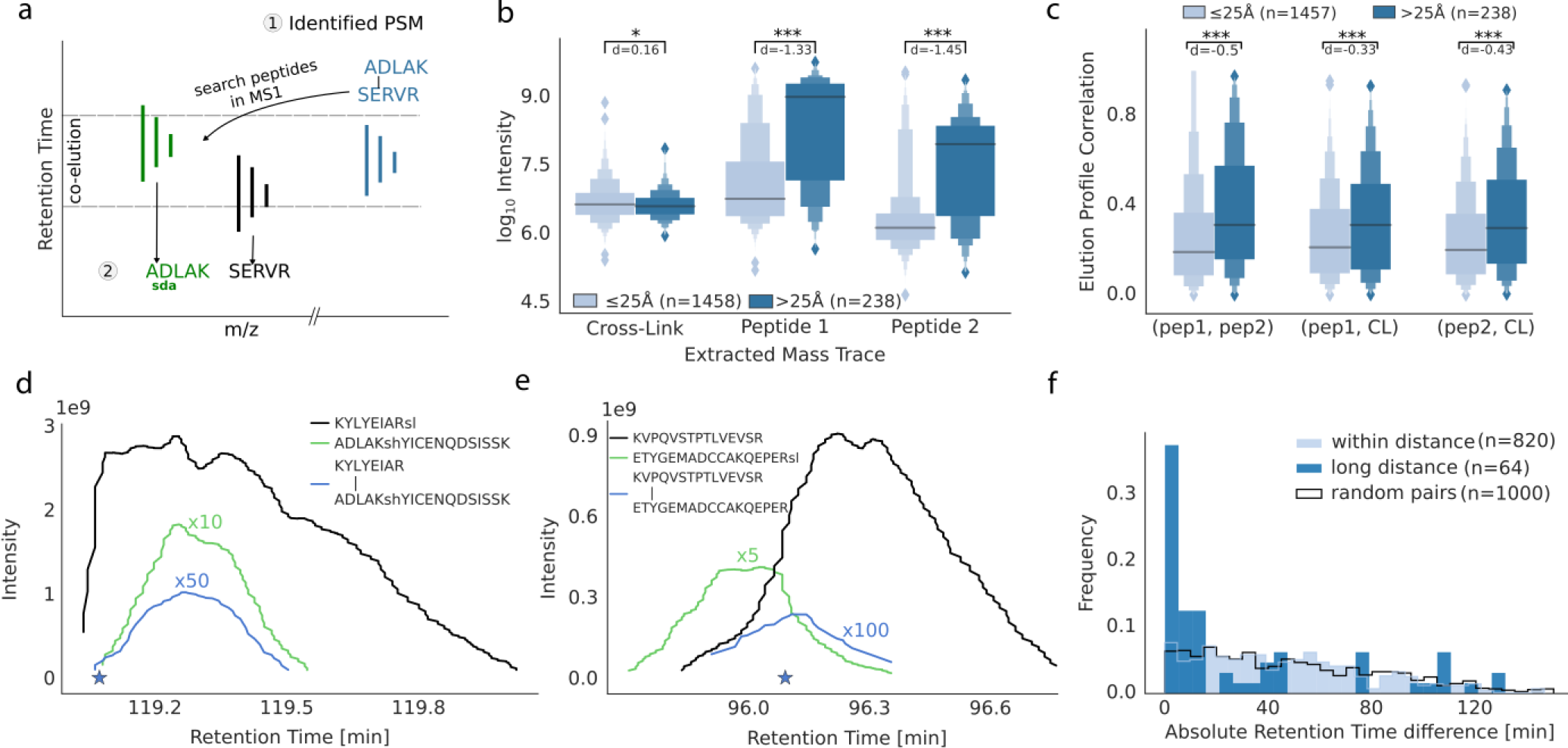
Analysis strategy and properties of LDL PSMs. (a) Non-covalent peptide search. Based on a crosslinked PSM (1), the individual peptide sequences are searched in the MS1 such that the summed mass equals the precursor mass of the identified crosslink (2). (b) Maximum intensity (along the m/z trace) for the identified crosslink and the m/z of the two individual peptides for links ≤ 25 Å and > 25 Å. (c) Spearman correlation of intensity profiles of the crosslink and the two individual peptides based on m/z matching in a RT window. (d-e) Examples of intensity profiles of two LDL. Filled stars mark the isolation time point of the precursor that yielded the identified crosslink. Scaling factors for lower intensity curves are written above the respective curves (e.g. x10 equals a factor of 10). Additional information about the PSMs can be retrieved through the uploaded results in PRIDE through the PSMIDs 7678478210 (d) and 7678602613 (e). (f) RT difference comparison of LDL and within-distance links.

We successfully extracted 1458 mass traces for PSMs of links within the distance cutoff and 238 mass traces for PSMs of LDLs. For these PSMs, we then compared the maximum intensity along the mass trace for the crosslink m/z and the two individual peptides m/z (Fig. 3b) within a window of ±90 seconds. Interestingly, the MS1 signals of long-distance links had significantly lower intensities than links fitting to the crystal structure, albeit with small effect size. In contrast, the MS1 intensities attributed to the individual peptides of LDL were higher by almost two orders of magnitude within the elution window compared to the control (peptides observed in crosslinks). This indicates a preference for co-elution of individual peptides with linked peptide pairs in the case of LDL but not within-distance links.

The high signal intensity of individual peptides of LDL around the elution of the LDL peptide made us wonder if they co-elute. We investigated the correlation of elution profiles more systematically by computing the Spearman correlation over the extracted ion chromatogram (XIC). While the absolute correlation is neither very high for the within-distance links nor for the LDL, the important feature is the difference between the two classes (Fig. 3c). The correlation of two single peptide m/z’s with each other - but also individually with the crosslinked m/z - are all significantly larger for the long-distance links compared to the within-distance links (p-value ≤ 0.001). The fact that the absolute value of the correlation is moderate is not surprising as it would be a precondition of non-covalent association that the individual peptides elute at an overlapping but not necessarily identical time, as is also seen from two examples of co-eluting and associating peptides (Fig. 3d-e). In the first example, all three m/z species start eluting at a similar time point. One of them is very abundant (MS1 intensity 1e9), reaching saturation and showing a long elution tail. This covers the complete elution time of the second peptide. As expected for an association product of the two, the LDL peptide then coincides with the elution of the second peptide. In a second example, the two individual peptides partially co-elute, and the LDL peptide is observed during the time of their overlapping elution.

To our surprise, some crosslinks that match the protein structure showed correlating MS1 intensities with their linear counterparts, despite a narrow matching time window. Retention on a reversed phase is usually very sensitive such that even peptide pairs with different crosslink sites show a different elution time.^26^ We therefore suspected the co-eluting MS1 intensities to be the baseline signal of our look-up strategy, which is solely based on m/z values and lacks confirmation through identification data. Hence, we checked for the RTs from the individual linear peptides relative to the crosslinks based on identifications instead of m/z matching alone. We compared the crosslink identifications with the closest RT from the linear identified peptides (with equal modifications and equal composition). The absolute difference of the individual RTs was mostly close to 0 minutes for the LDL PSMs and approximately uniformly distributed for within-distance PSMs (Fig. 3f). The added control (random pairings of RTs from linear identified peptides that were also part of a crosslinked peptide) closely resembles the within-distance PSM distribution. However, only 50% of the PSMs have a RT difference smaller than 10 minutes. The remaining PSMs have a large RT difference which reduces the possibility of co-elution. Interestingly, PSMs with a RT difference smaller than 10 minutes have an average score of 10.0 (n=32), while the remaining PSMs (n=32) have an average score of 6.7. Possibly, the lower score indicates imprecise peptide identifications and thus wrong RT times. In addition, matches with large RT differences can still originate from wrong identifications. Like target-decoy matches in a crosslink, in a NAP one of the peptides could be correct and the other might be a random match. In these cases, the RT difference would also be randomly distributed.

#### In-source fragmentation reduced the number of non-covalently associated peptides

Based on the results above, one would predict NAPs to form even without prior crosslinking. The phenomenon should only depend on peptide concentration and their affinities. We therefore investigated a four-protein mix without any cross-linker addition and wondered if non-covalent associated peptides could be identified. Note that here we changed to a Q Exactive High-field. Indeed, we identified 24 non-covalent peptide associations (Fig. 4). The formation of NAP is thus also observable in linear proteomics that do not involve any cross-linking chemistry. However, the number of NAP identifications is low and unlikely to affect linear proteomics.

**Figure 4:**
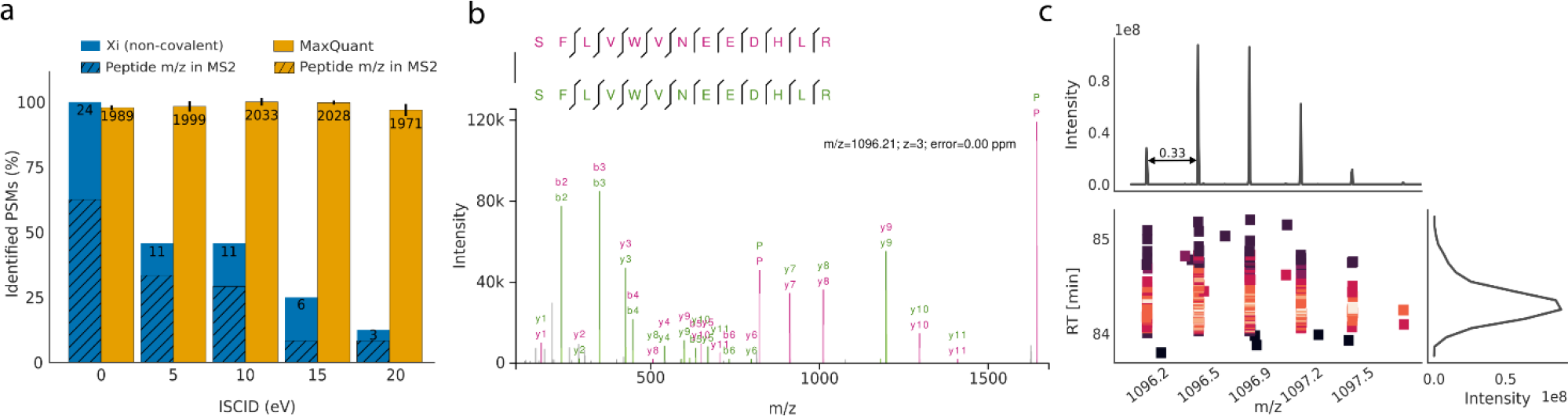
Non-covalently associated peptide identifications in non-crosslinked samples. (a) Number of PSMs after 5% PSM FDR in a non-covalent search and linear identifications (1% FDR). Peptide m/z fraction refers to occurrences where the individual peptide or precursor peaks are found in multiple charge states in the MS2 spectrum. (b) Non-covalent peptide identification with charge state 3, individual peptide peaks (P) were identified with charge 2 (822.41 m/z) and charge 1 (1643.82 m/z). (c) MS1-derived peptide feature for the PSM displayed in (b). Top panel shows the summed intensity over the m/z bins. Bottom panel shows the m/z over the RT color-coded by the intensity. Right panel shows the summed intensity over the RT.

Since the involved forces leading to an interaction are expected to be rather weak, employing in-source collision induced dissociation (ISCID) should reduce the number of identified NAP. Using an ISCID of 0, 5, 10, 15 and 20 eV, we find 24, 11, 11, 6 and 3 NAP identifications at 5% PSM FDR (Fig. 4a). Increasing the ISCID from 0 to 20 results in a 90% decrease of NAPs identifications. As a control, we also investigated how linear peptide identifications were affected by these voltages for ISCID and observed only a minor detrimental effect. Predominantly, we saw self-associations of the same peptide with all ISCID settings (88%, 64%, 73%, 33% and 67%) for 0, 5, 10, 15 and 20 eV ISCID). Also, in crosslinked HSA we saw many self-links of peptides, which initially perplexed us as these would indicate protein dimerization despite us having isolated and analyzed the monomer. These crosslinked peptides now pose strong candidates for NAPs as well. This indicates that special care must be taken when homo-multimers are investigated via CLMS. Note, that homo-multimers are not necessarily identified through crosslinks of the same peptide in both instances of the protein. Crosslinks involving overlapping peptide sequences can also indicate homo-multimerization (see Fig. S5).

We noticed a feature of MS2 spectra of NAPs that may help identifying them in future. The intact peptide peaks in multiple charge states up to the NAP’s precursor charge state are frequently observed and are of high intensity (Fig. 4a and b). We encountered this in 62% of cases for the ISCID data set of 0 eV. We are unaware of such charge reduced precursor ions in HCD fragmentation spectra of linear peptides and do not see a single occurrence in our linear peptide data. This adds to NAPs being revealed at MS1 level through their overlapping elution with the individual linear peptides. It is unclear if NAP can be avoided altogether. However, critical assessment of the ionization settings appears to be advisable for CLMS analyses.

For the analysis of proteins via native MS, one should be aware that these unspecific associations might be possible too, even under ‘normal’ LC conditions as we have used here. The exact conditions that support the formation of NAPs are not known.^29^ However, previous studies found that electrostatic interactions lead to increased stability of non-covalent complexes^30,31^, but also solvent composition and ionization settings^29,32^ are crucial. Likely, any parameter influencing the ionization such as instrument architecture and flow rates play a role. We therefore tested the influence of three flow rates on the formation of NAPs but found no differences within our experimental set-up (Fig. S3).

For crosslinking mass spectrometry experiments, NAPs pose a challenge. Crosslinking experiments using SDA or similar reagents are more susceptible to NAP identifications since the crosslinker can form loop-links on lysine residues, resulting in the same modification mass as a crosslinked peptide pair. However, the formation of NAPs does not depend on the crosslinker since we also observed their formation in non-crosslinked samples. Therefore, in theory, other crosslinkers will also lead to NAPs. A critical assessment of the specific instrument ionization settings is thus crucial for successful analysis of CLMS experiments. If the possible presence of NAPs is ignored, they will lead to wrong distance constraints. Even though structural modeling approaches are to some extent robust to the number of false positives^27^ the influence of a systematic source of false positives is unknown. Experiments that aim to reconstruct the rough topology of protein complexes are at high risk to draw false conclusions from these false ‘crosslinks’. Wrong inter-protein links and wrong intra-protein / loop-links might lead to inconclusive results. Therefore, we strongly suggest reducing the possibility of NAPs, either by optimizing acquisition settings or heuristic post-acquisition filters.

#### Significance of non-covalently associated peptides

We observe NAPs here during the analysis of an SDA-crosslinked protein. While SDA is of central importance to high-density CLMS and the development of crosslinking for protein structure determination, this is a very young research area with currently few followers. Nevertheless, NAPs do not require the presence of SDA as we show by our analysis of a standard four-protein mix, without any crosslinking. The possible impact of NAPs goes into several directions, where few NAPs could make an impact. Self-association of loop-linked peptides would also occur with crosslinkers such as BS3 or DSS, leading to the possibility of misidentifying NAPs as cross-links. This would then lead to a false biological conclusion, namely that a protein self-associates to form homodimers. Cleavable crosslinkers have the advantage that if a full set of signature peaks is observed, NAP formation can be ruled out. Unfortunately, the set of signature peaks is not always complete.^33^ Secondly, our analysis showed that NAPs yield excellent spectra, often better than crosslinked peptides. When not considering NAPs, these good spectra can match only one of the associated peptides correctly, while for the second one the mass would be off by the assumed presence of a crosslink. This can only lead to a false target-target (TT) hit or target-decoy (TD) hit. Indeed, we found in our analysis an example (Fig. 5) where a high scoring TD from a crosslink search matched a TT during a NAP search with improved confidence. In routine analyses of protein complexes relatively few crosslinks are being detected so few high-scoring TDs may noticeably reduce the identified links. This was not the case in our analysis but should not be dismissed outright and warrants further attention. Finally, the presence of biologically not functional peptide-peptide complexes in the gas phase suggests that also during the analysis of much larger proteins with many more interaction possibilities may lead to such non-biological associations. Consequently, Native mass spectrometry may require the development of appropriate controls as has been suggested before^4^.

## Conclusion

Self-associations of peptides in solution has been shown to yield stable oligomers that endure the ionization process^32^. In addition, the preservation of non-covalent associations throughout ESI is exploited by native mass spectrometry. Here, we show that peptides with very similar chromatographic RT behavior can also remain together during the ionization process under normal liquid chromatography conditions as they are used in bottom-up proteomics. This implies that the association process can be unspecific and occur during normal LC-MS analysis. At the very least, the CLMS field should be aware of this. Pointing at ionization parameters and post-acquisition tests, we hope to assist the field in spotting and counteracting this effect.

**Figure 5:**
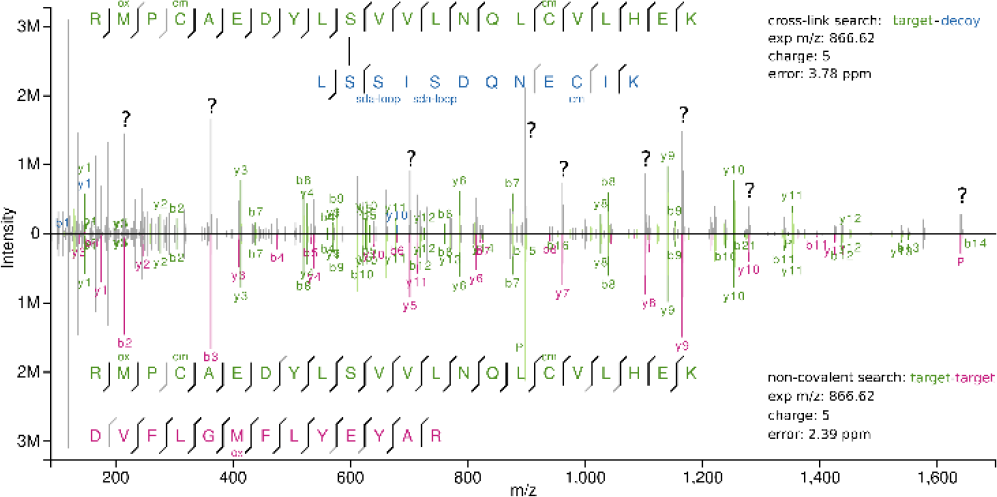
Butterfly plot of the same spectrum with different possible explanations. Upper panel shows the annotation from a cross-linking search (target-decoy identification). Lower panel shows the annotation from a non-covalent search (target-target identification). Q Exactive acquisition; raw file: *V127_K*; scan: 50038.

## Associated Content

### Supporting Information

Conceptual drawings of crosslinks and non-covalently associated peptides, results using cleavable crosslinking search software and results from acquisitions with different flow rates.

The mass spectrometry raw files, peak lists and search engine results have been deposited to the ProteomeXchange Consortium (http://proteomecentral.proteomexchange.org) via the PRIDE partner repository^25^ with the dataset identifier PXD010895. In addition, the PSMs at 5% FDR are available online using xiVIEW^34,35^:

- Velos, HSA data (https://xiview.org/xi3/network.php?upload=34-08362-96692-34003-27750)
- QE, HSA data (https://xiview.org/xi3/network.php?upload=35-49786-17881-94522-79322)
- Q Exactive HF, protein mix (https://spectrumviewer.org/viewSpectrum.php?db=ISCID)

## Author Information

### Corresponding Author

*

### Author Contributions

The manuscript was written through contributions of all authors.

## Acknowledgment

This work was supported by the Einstein Foundation, the DFG [RA 2365/4–1, 25065445], and the Wellcome Trust through a Senior Research Fellowship to JR [103139] and a multi-user equipment grant [108504]. The Wellcome Centre for Cell Biology is supported by core funding from the Wellcome Trust [203149].

---

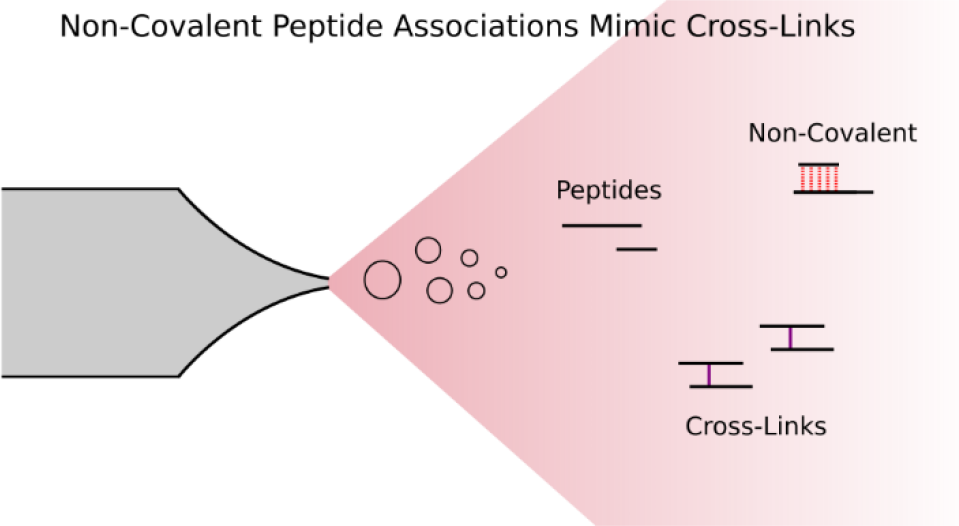
Insert Table of Contents artwork here.

