## Supporting Information for "Non-covalently-associated peptides are observed during liquid chromatography-mass spectrometry and affect crosslink analyses"

#### Contents

**S1 Visualization of crosslinks and non-covalently associated peptides** **2**

**S2 CLMS identifications assuming cleavability of SDA** **2**

**S3 Flow Rate Analysis on Q Exactive High-field** **3**

**S4 Falsely identified crosslink suggesting homo-dimerization** **4**

*

### S1 Visualization of crosslinks and non-covalently associated peptides

Fig. S1 visually describes the different types of peptides relevant for the main manuscript. Importantly, only the crosslinked peptide (a) and the non-covalently associated peptide (c) have the same mass because of the loop-link in c). This mass ambiguity is the reason that non-covalently associated peptides can be misidentified as crosslinks. More details on general crosslinking nomenclature can for example be found in *Rappsilber* [1]. Note that the two peptides in c) do not need to have the same sequence.

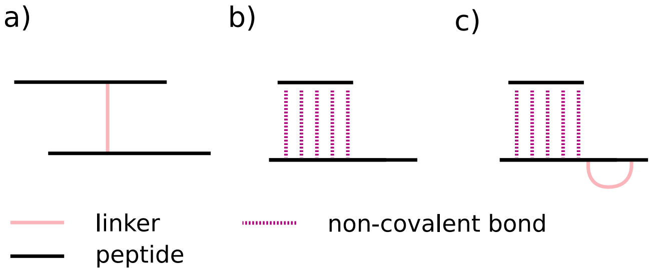

Figure S1: Visualization of different peptide definitions. (a) a typical crosslink between two peptides. (b) a non-covalent peptide association between two peptides. (c) same as (b) but one of the peptides is loop-linked, the mass of this species is the same as that of a crosslinked peptide.

### S2 CLMS identifications assuming cleavability of SDA

In this section we used MeroX [2] to identify MS-cleavable crosslinker products from the Q Exactive (QE) data set. We used the same settings for MeroX (v. 1.6.6.6) as in [3]. The SDA reaction is assumed to be cleavable when involving a carboxylic acid functional group [3]. The individual search results were combined using the MeroX Merger (v. 1.2) with the -P 5 setting to set the desired FDR cut-off to 5%. From the merged results we extracted the unique links and computed their distance in the crystal structure of HSA (PDB: 1AO6). For the QE data, 184 unique links were identified of which 160 could be mapped to the crystal structure. 38% (61 links) were long-distance links ($C_\alpha distance \geq 25$Å), while 62% (99 links) matched the distance constraint. For the Velos data, 34 unique links were identified of which 29 could be mapped to the crystal structure. 21% (6 links) were long-distance links, while 79% (23 links) matched the distance constraint. The results are consistent with the presented data in Figure 1 of the manuscript. The distance histogram for the QE data shows a very prominent enrichment of false positives exceeding the distance threshold. While in both cases the desired FDR is not met, we hypothesize that the Velos results are suffering from the low number of identified links. Therefore, reliable FDR estimation is hindered.

In general, Fig. S2a-b shows that MeroX is also able to identify the non-covalent peptide associations using a cleavable crosslinker search. However, it is difficult to judge how many of the identified crosslinks below the 25 Å cut-off are true crosslinks (assuming cleavage of the crosslinker) and how many are non-covalent associations. Since the search itself is not aware of any distance constraint, an obvious assumption is that non-covalent associations should be distributed without preference below and above the distance cut-off. In contrast, true crosslinks will be enriched below the distance cut-off. Visually projecting the number of long-distance links to the area below the distance cut-off indicates that a large portion of the within-distance links are in fact non-covalent associations.

In addition, we also analysed the retention times (RT) from linear peptides with SDA modifications (e.g. loop-linked or hydrolyzed crosslinker, see [4] for visualizations of the modifications). Interestingly, the RT of linear peptides is approximately increased by 24 minutes with a single sda-loop modification (Fig. S2c). Subsequently, the RT is almost doubled (42 minutes) when two loop-links were found in a peptide compared to the unmodified version. This information can be used to compare the RT of the linear peptides that were identified in a crosslink / non-covalent peptide association. We used the simplified assumption that identifications are true crosslinks when the distance constraints were met and non-covalent association otherwise. In Fig. S2d, the RT difference between the two peptides in a crosslink / non-covalent association is shown. Initially, we tried to map the individual peptide sequences identified by MeroX to the linear (modified) peptide identifications from MaxQuant. For this one of the two peptides identified in a crosslink by MeroX was assumed to carry a loop-link modification. Under these assumptions only a small number of crosslinked peptides yield a RT for both peptides. The reason is that the individual peptides identified by MeroX were not identified in their loop-linked form in MaxQuant. For peptides that are crosslinked, the RT difference from the individual peptides should be randomly distributed. For peptides that are non-covalently associated, the RT difference from the individual peptides should be closely distributed zero. Because MeroX does not search for loop-link modifications in the search for non-covalently associated peptides the RT difference that is introduced through this modification needs to be accounted for. Therefore, the expected RT difference for the individual peptides from non-covalent associations is on average 24 minutes. Indeed, the two RT difference distributions from crosslinks and non-covalent associations look different and match the above described expectation (Fig. 3c). However, the large enrichment of within-distance links with a very small RT difference hints on these identifications being non-covalent peptide associations.

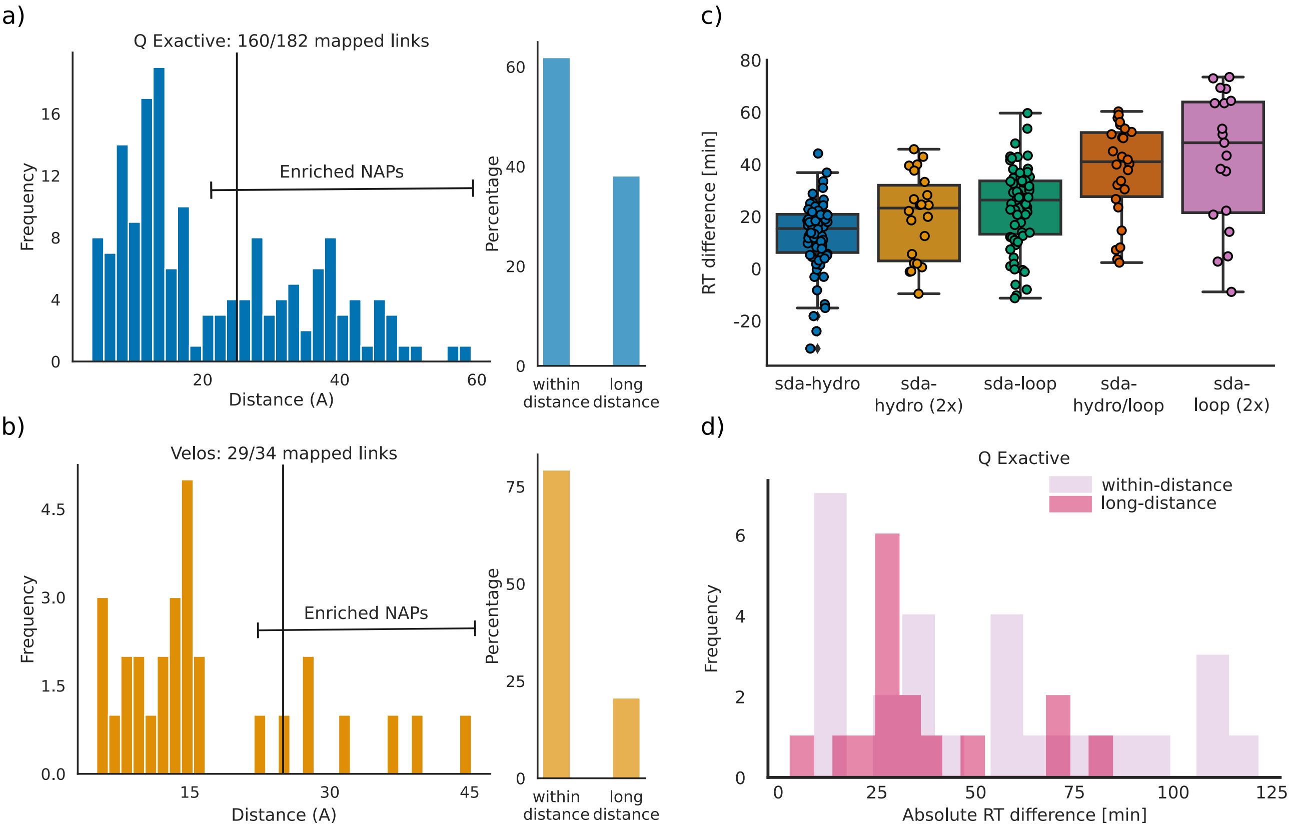

Figure S2: MeroX results and MaxQuant results. (a) Distance-histogram of unique links identified with MeroX for Q Exactive data. (b) Distance-histogram of unique links identified with MeroX for Velos data. (c) Retention time difference of unmodified and crosslinker modified linear peptides identified with MaxQuant. (d) RT mapping of PSMs from a) to linear identifications (MaxQuant search) without RT adjustment for modifications. *Note*: RT - retention time, NAP - non-covalent association, sda-hydro and sda-loop refer to modified crosslinkers [4].

### S3 Flow Rate Analysis on Q Exactive High-field

To further investigate the effect of different flow rates on the formation of non-covalent associations we acquired the protein mix (non-crosslinked sample) on the Q Exactive High-field with three flow rates (in triplicates): 0.2 $\frac{\mu L}{min}$, 0.25 $\frac{\mu L}{min}$ and 0.3 $\frac{\mu L}{min}$. The differences in the number of identifications were only small (Fig. S3a) and comparable to the results from the main text (24 PSMs with IS-CID 0). To achieve the desired FDR cut-off of 5% the results were cut after the first decoy hit (S3b).

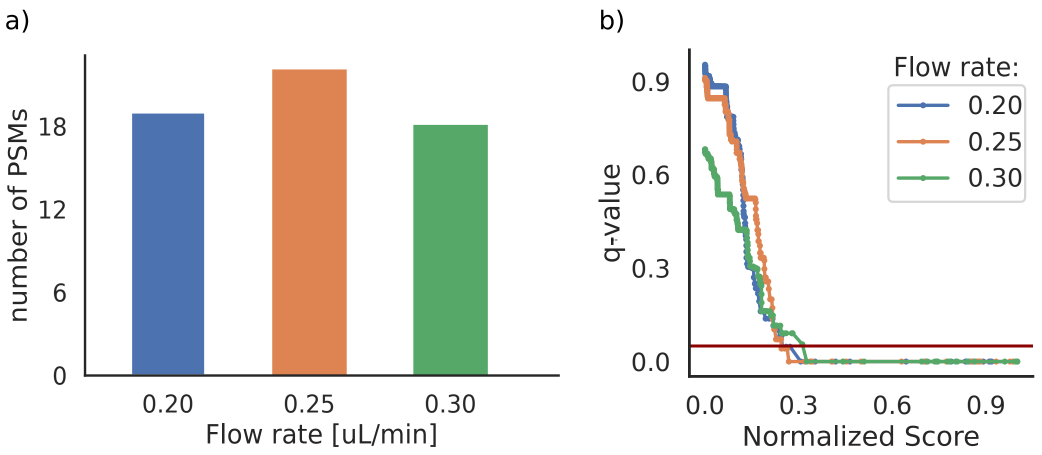

Figure S3: Flow rate analysis. a) shows the number of PSMS passing a 5% FDR cut-off (or until the first decoy hit if the cut-off is exceeded). b) shows the q-value trend with decreasing scores. The horizontal red line marks the 5% cut-off. The search engine score was normalized by division through the respective maxima.

#### S4 Falsely identified crosslink suggesting homo-dimerization

The spectrum in Fig. S4 shows an example of a crosslink that can falsely lead to the assumption of homo-dimerization.

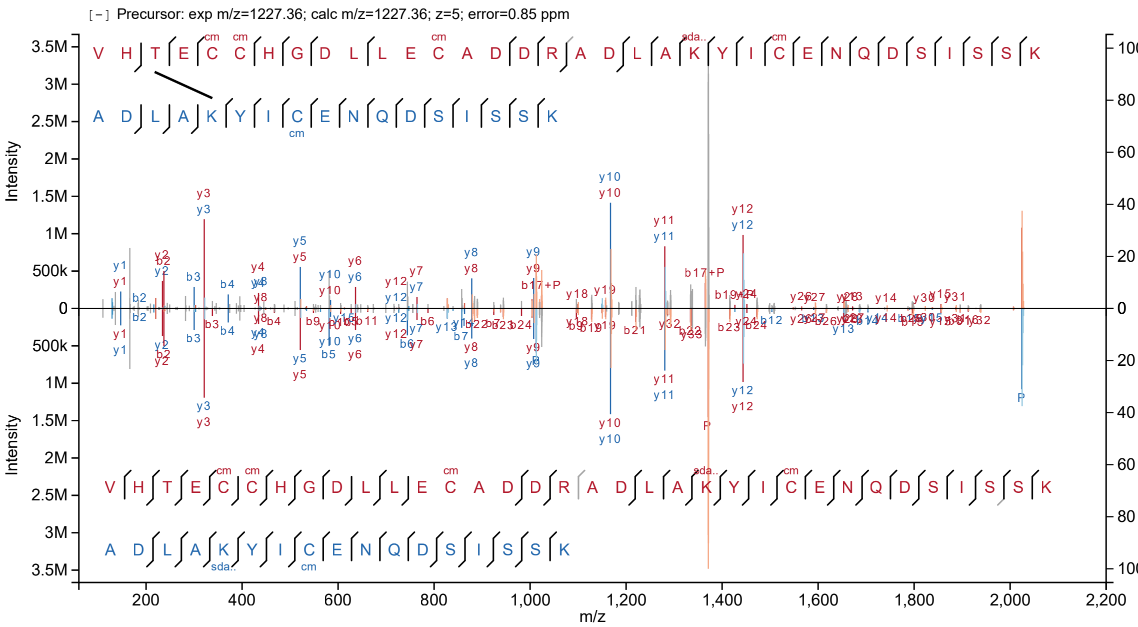

Figure S4: Alternative explanation for a crosslink that would suggest homodimerization. Upper panel, annotation from non-covalent search. Lower panel, annotation from crosslink search. Raw file: V127_J; scan: 34926
